# Synaptic engram underlies memory

**DOI:** 10.64898/2026.07.30.740767

**Authors:** Dae Hee Han, Hoonwon Lee, Jongrok Do, Pojeong Park, Hyunmin Park, Hyungji Lee, Yeonjun Kim, Chang-Ho Kim, Jooyoung Kim, Chaery Lee, Jun-Yeong Baek, Myunghyun Cheon, Ji-il Kim, Dong Il Choi, Doyun Lee, Bong-Kiun Kaang

**Author notes:** These authors contributed equally (Dae Hee Han, Hoonwon Lee, Jongrok Do).

## Abstract

Memory encoding cells, or engram cells, are sparsely distributed across the brain. Recent findings have shown that memory formation coincides with the strengthening of interregional synaptic connectivity between engram ensembles. However, whether this learning-induced synaptic reorganization causally supports memory, independently or in concert with heterosynaptic and intrinsic plasticity, remains elusive. Here we developed optically implemented spike-timing-dependent plasticity (oSTDP), which induces synaptic depression selectively at target synapses by controlling the relative timing of pre- and postsynaptic activity. In vivo application of oSTDP to the synapses between entorhinal cortex (EC) and dentate gyrus (DG) engram cells caused a significant synaptic depression and the resultant memory impairment. Our results demonstrate that plasticity of functional connectivity between memory-encoding ensembles causally underlies memory.

## Main Text

Experience forms memory which serves as a building block of the internal model by which external stimuli drive various behavioral outputs. Memory is believed to exist in the brain as a physical substrate, or an engram (1). The activity of a sparse set of neurons activated during learning, or engram cells, is necessary for memory retrieval and sufficient to recall memory in the absence of a retrieval cue, constituting the cellular substrate of an engram (2–5).

Spatiotemporal summation of synaptic inputs drives diverse patterns of neural firing that recapitulate the neural representations for cognitive functions including memory (6, 7). Therefore, plasticity occurring at learning-relevant synaptic connections is widely considered as a mechanism of episodic memory and long-term potentiation (LTP) of synapses has been regarded as its primary cellular mechanism (8–10).

Then what are the learning-relevant synapses? Given the central role of engram cells in episodic memory, learning-induced synaptic reorganization in these cells has emerged as a potential mechanism underlying memory storage (11–14). Particularly, synaptic inputs to postsynaptic engram cells arriving from presynaptic engram cells, hereafter referred to as engram synapses (15–17), undergo synaptic strengthening during memory formation, suggesting that these are the synaptic loci where learning-relevant plasticity occurs.

Given the sparse nature of pre- and postsynaptic engram cells, however, engram synapses constitute a minute proportion of the entire synaptic connections. Whether memory representation can depend on plasticity at such a small synaptic population remains unclear. If so, selectively reversing learning-induced changes in these synapses will impair memory expression. Alternatively, homeostatic plasticity of non-engram synapses and cell intrinsic plasticity in excitability may complementarily underlie memory representation (18, 19).

However, testing whether synaptic potentiation of engram synapses causally serves memory by selective manipulation is challenging because these synapses are small and spatially intermingled with non-engram synapses. Here, by leveraging optically-implemented spike-timing dependent plasticity (oSTDP), we show that the selective disruption of strengthened functional connectivity between interregional engram ensembles impairs memory.

## Results

### Enhanced strength of engram synapses in the perforant paths following contextual fear memory formation

Beyond merely correlating with learning, the activity of DG engram cells is required for the consolidation of contextual fear memory (20, 21). The medial and lateral entorhinal cortices (MEC/LEC), major input regions of DG, provide enhanced excitatory synaptic input to DG engram cells (22–25). However, which cellular population in the EC provides this increased synaptic input following learning remains unknown.

To examine whether synaptic connections between EC and DG engram cells are strengthened following contextual fear conditioning, we labeled EC and DG engram cells using a cFos-based Tet-on system in combination with a TRAP system (Arc-CreER^T2^; fig. S1 and S2) (26), and applied the previously established synapse labeling technique (dual-eGRASP)(15) to visualize their connectivity (fig. S1, A and B; Fig. 1, A to C). DG engram dendrites were labeled with mScarlet, while sparse sets of non-engram dendrites were labeled with iRFP (Fig. 1C).

**Fig. 1.**
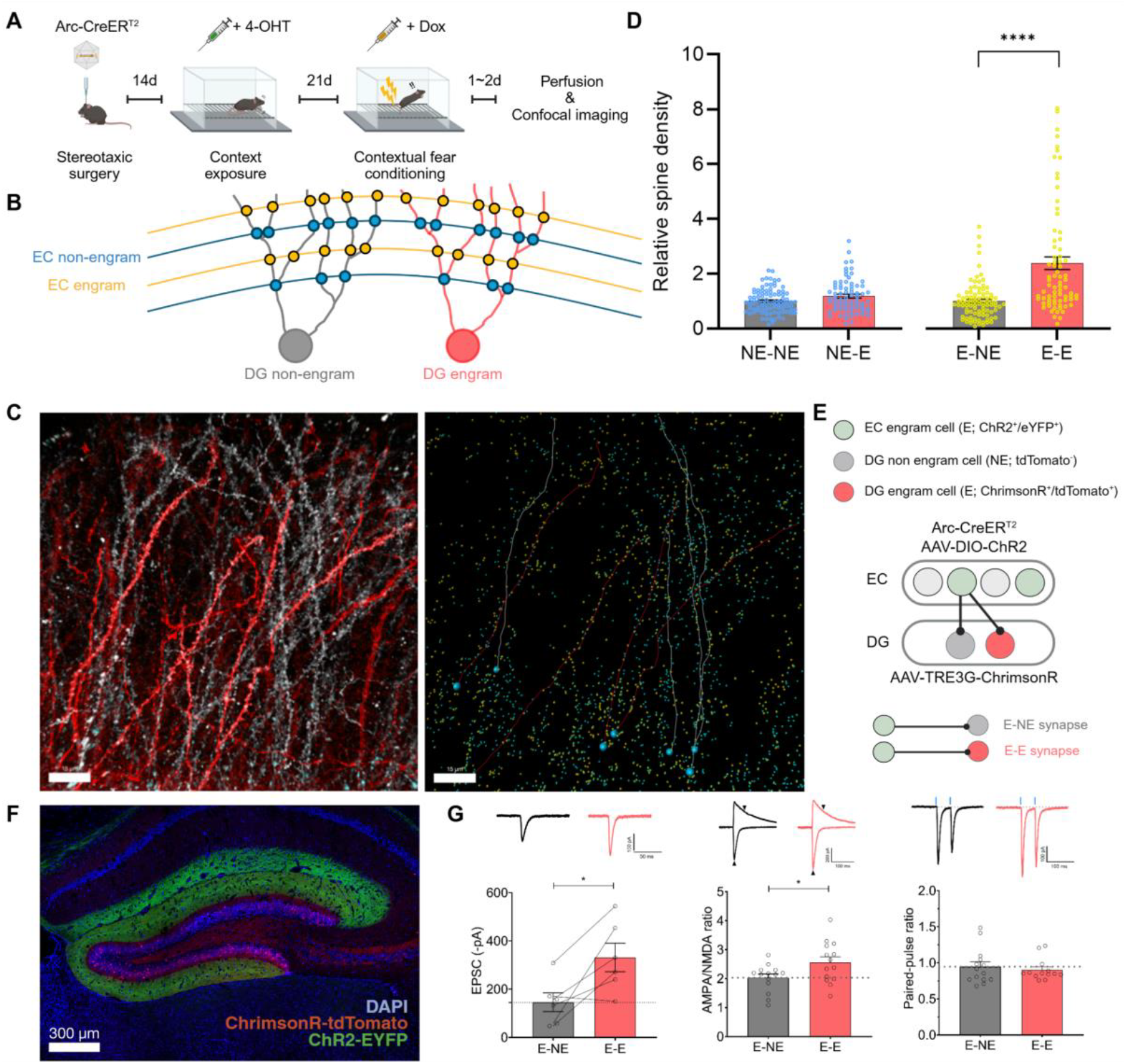
Enhanced functional connectivity between EC and DG engram cells following learning. **(A)** Schematics showing experimental schedules for dual-eGRASP imaging. **(B)** Schematics illustrating synaptic and dendritic identities in dual-eGRASP experiments. **(C)** Representative images showing dual-eGRASP signals. Left, postsynaptic engram or non-engram dendrites with cyan GRASPs (non-engram synaptic inputs) and yellow GRASPs (engram synaptic inputs) signals. Right, distribution of cyan GRASPs and yellow GRASPs. Scale bar: 15 μm. **(D)** Relative spine densities (see the method) of each type of synapses defined by pre-/postsynaptic identities (NE-NE *n* = 91, light green; NE-E *n* = 75, blue; E-NE *n* = 91, tangerine; E-E *n* = 75, red; *\*\*\*\*P < 0*.*0001*, Mann Whitney test). **(E)** Experimental schematics for AMPA/NMDA ratios and PPRs measurements in DG engram and non-engram cells, with EC engram cell inputs being stimulated. **(F)** Representative image showing hippocampal slices with EC engram inputs labeled with ChR2-EYFP and DG engram cells labeled with ChrimsonR-tdTomato. Scale bar: 300 μm. **(G)** Left, optically-evoked EPSCs (oEPSCs) amplitudes of E and NE pairs by MEC engram inputs (*n* = 6 pairs; \**P <* 0.05, paired *t*-test). Middle, AMPA/NMDA ratios of DG non-engram and engram cells to synaptic inputs from MEC/LEC engram cells (E-NE *n* = 14, E-E *n* = 14 cells; *\*P <* 0.05, unpaired *t*-test). Right, PPRs of DG non-engram and engram cells to synaptic inputs from MEC/LEC engram cells (inter-stimulus interval 50 ms; E-NE n = 14, E-E n = 13 cells; *P* = 0.4794, unpaired *t*-test). Inset shows corresponding representative traces of oEPSCs, AMPA/NMDA ratios or PPRs of non-engram cell (black) and engram cell (red).

Simultaneously, yellow and cyan GRASP signals marked synaptic contacts formed by EC engram and non-engram cells, respectively. Yellow GRASP synapses were significantly denser on mScarlet+ dendrites (E-E) than on iRFP+ dendrites (E-NE; Fig. 1D), whereas cyan GRASP synapse density exhibited no significant difference across dendrite identities (NE-E vs. NE-NE; Fig. 1D). These results indicate that EC and DG engram cells are preferentially connected following contextual fear conditioning.

Next, DG and EC engram cells were labeled with ChrimsonR-tdTomato and ChR2-EYFP, respectively (fig. S1, C and D; Fig. 1E) to identify DG engram cells with tdTomato while EC engram axons were stimulated with 473 nm laser pulses in transverse hippocampal slices (Fig. 1,E and F). DG engram cells (tdTomato+; E) received stronger synaptic inputs from EC engram cells compared to DG non-engram cells (tdTomato-; NE), as evidenced by larger optically-evoked excitatory postsynaptic currents (oEPSCs) and a higher AMPA/NMDA ratio (Fig. 1G). Paired-pulse ratios (PPRs) of EC engram inputs did not show a significant difference between DG engram and non-engram cells (Fig. 1G). Together, these data indicate that EC engram cells exert stronger synaptic drive to DG engram cells than to non-engram cells, mainly by enhanced postsynaptic AMPA receptor-mediated synaptic transmission and greater connectivity.

### Optically-implemented spike-timing-dependent plasticity (oSTDP)

To manipulate engram synapses selectively, we devised a method based on spike-timing-dependent plasticity (STDP) that induces synaptic depression at synapses between EC-DG engram cells without affecting spatially-intermingled non-engram synapses. In STDP (27–29), the precise temporal difference between pre- and postsynaptic firing determines the magnitude and direction of the resulting synaptic plasticity, and we reasoned that this property could enable targeted manipulation when applied to engram synapses (Fig. 2, A to D).

**Fig. 2.**
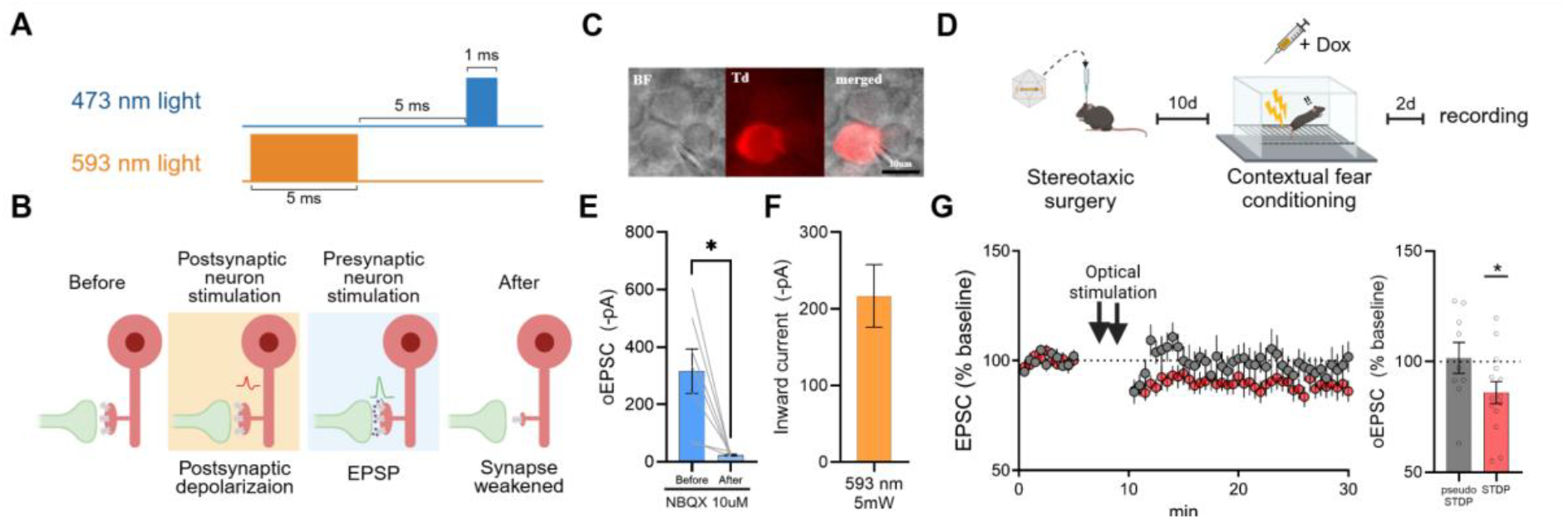
Optically-implemented spike-timing-dependent plasticity. **(A)** Schematics for the description of oSTDP stimulation protocol used for *in vitro* experiments. **(B)** A rationale for implementing optical STDP (oSTDP) for specific weakening of engram synapses. **(C)** A representative image showing an engram cell (tdTomato+) identified in an acute hippocampal slice. BF, bright field; Td, tdTomato. Scale bar: 10 µm. **(D)** Schematics showing experimental schedules for slice electrophysiology. **(E)** MEC input evoked oEPSCs are blocked by NBQX (10 µM; *n* = 7 cells, * *P* < 0.05, Wilcoxon test). **(F)** Inward current of ChrimsonR+ neurons in response to 593 nm (5 mW, 5 ms) light stimulation (*n* = 13 cells). **(G)** A time course of MEC input evoked oEPSCs in DG engram cells during baseline and following either oSTDP or pseudo oSTDP stimulation (grey, pseudo oSTDP, *n* = 9 cells, *ns*; red, oSTDP, *n* = 14 cells, * *P < 0*.*05*; one sample *t*-test with theoretical mean = 100).

To test this idea, we first characterized the STDP rules in the MEC-DG projection (fig. S3, A to D) (30, 31). Significant synaptic depression was observed when postsynaptic depolarization preceded presynaptic stimulation by a few milliseconds, and prior induction of synaptic potentiation changed the plasticity kernel toward synaptic depression (fig. S3, B and C).

We then tested whether STDP could be implemented optically in a synapse-specific manner *in vitro*. We expressed ChR2-EYFP in a large excitatory neuronal population in the MEC and ChrimsonR-tdTomato in DG engram cells (fig. S3E). Given the distinct excitation spectra of ChR2 and ChrimsonR (Fig. 2, E and F) (32), we reasoned that alternating two wavelengths (473 and 593 nm) separated by a few milliseconds could reproduce the activity pattern that induces STDP (oSTDP; Fig. 2A). Repeated pairing of postsynaptic depolarization followed by presynaptic release significantly decreased the amplitude of oEPSCs, indicating successful depotentiation of MEC synaptic input to DG engram cells by oSTDP (Fig. 2G). This reduction was absent when the same pairing was delivered with a long temporal delay (250 ms; pseudo oSTDP; Fig. 2G).

### Weakening of functional connectivity in EC-DG engram ensemble by oSTDP

We next expressed ChR2-EYFP in EC engram cells to selectively manipulate the EC-DG engram connectivity using oSTDP. This stimulation reduced the amplitude of oEPSCs measured in DG engram cells (Fig. 3, A to C). The effect was strictly dependent on paired activation, as omitting either 473 nm or 593 nm light yielded no change in oEPSCs (Fig. 3C). Furthermore, oSTDP stimulation did not alter EPSC amplitudes in DG non-engram (ChrimsonR-) cells (fig. S4, A to C), supporting the idea that oSTDP-dependent synaptic depression is restricted to engram synapses.

**Fig. 3.**
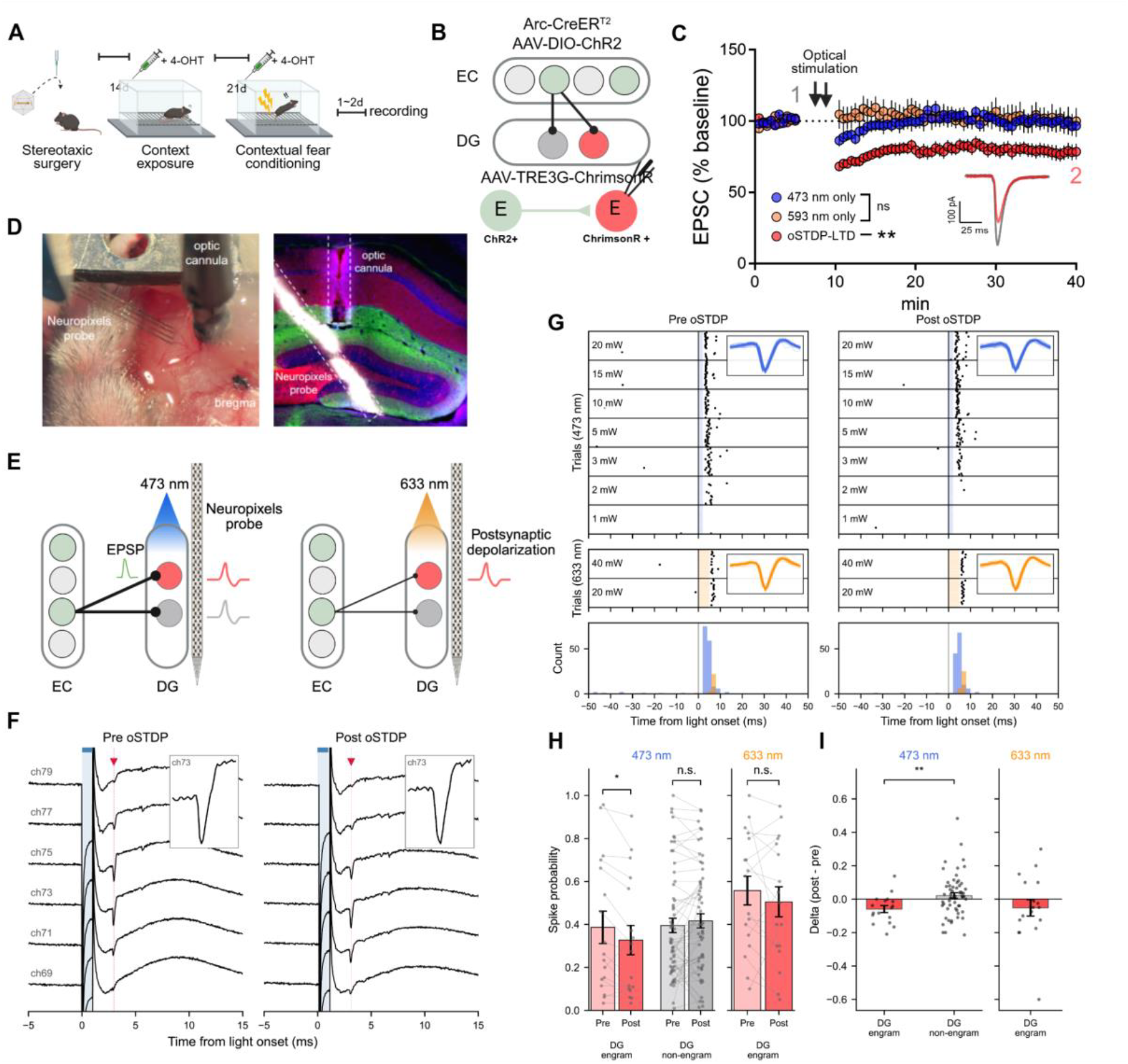
Weakening of functional connectivity of EC-DG engram ensembles by oSTDP-LTD. **(A)** Schematics showing behavioral schedule for *in vitro* oSTDP-LTD experiments in engram synapses. **(B)** Experimental scheme for *in vitro* validation of oSTDP-LTD protocols. **(C)** Time course of EC engram input-evoked oEPSCs amplitude of engram synapses before/after each type of optical stimulation (red, oSTDP-LTD, *n* = 12 cells, \*\**P<*0.01; blue, 473 nm lights only, *n* = 6 cells, *P =* 0.4741; tangerine, 593 nm lights only, *n* = 6 cells, *P =* 0.8176; paired *t*-test with baseline). Insets show merged representative traces obtained at the designated time point of oSTDP-LTD trace (grey, 1; red, 2). Statistic tests were conducted by comparing average amplitude of the last 5 sweeps of EPSCs during baseline period and the last 5 sweeps of EPSCs 30 min after optical stimulation of each type. **(D)** Left, image showing the Neuropixels and optic-cannula preparation. Right,a representative image displaying axonal projections from EC engram cells (ChR2-EYFP), DG engram cells (ChrimsonR-tdTomato), DAPI, and Neuropixels probe track (DiD). **(E)** Experimental schematics for Neuropixels recording with dual-opsin optogenetics. **(F)** Raw voltage traces from an example unit around 473 nm light stimulation during the Pre-oSTDP (left) and Post-oSTDP (right) optogenetic stimulation phases; blue bar, light-on window; red dashes, example spikes evoked by the 473 nm light pulse. **(G)** Raster (top) and peristimulus-time histograms (bottom) for a representative unit during optogenetic stimulation phases before and after oSTDP; insets show individual waveforms of detected spikes (blue, 473 nm light stimulation-evoked; orange, 633 nm light stimulation-evoked). **(H)** Left, spike probability during the 0–10 ms window after 473 nm light onset in the Pre- and Post-oSTDP optogenetic stimulation phases for DG engram (n = 17, \**P* < 0.05, Wilcoxon signed-rank) and non-engram (n = 58, *ns*; Wilcoxon signed-rank) units. Right, spike probability after 633 nm light onset for DG engram (n = 17, *ns*; Wilcoxon signed-rank). **(I)** oSTDP-induced changes (Post - Pre) in 473 nm-evoked spike probability (left) for DG engram (n = 17) and non-engram (n = 58) units (Mann–Whitney U \*\**P* < 0.01) and in 633 nm-evoked spike probability (right) for DG engram (n = 17). Data are mean ± s.e.m. *ns*, not significant.

This synaptic weakening is primarily mediated by the postsynaptic removal of functional AMPARs, as evidenced by a decreased AMPA/NMDA ratio after oSTDP-LTD stimulation in engram synapses *in vitro* (Fig. 1G; fig. S5, B to D). Furthermore, the PPRs remained unchanged, indicating that oSTDP did not affect the presynaptic release properties of engram synapses *in vitro* (Fig. 1G and fig. S5E).

To test if oSTDP-LTD could selectively disrupt EC-DG engram functional connectivity *in vivo*, we developed a novel approach combining dual-opsin optogenetics (ChR2 and ChrimsonR) with Neuropixels recording. By stimulating ChR2-expressing EC axons in the DG and ChrimsonR-expressing DG neurons with spectrally distinct light, we assessed two properties of the recorded DG units (Fig. 3, D and E). First, based on their responses to ChrimsonR stimulation, recorded units were classified as ChrimsonR-expressing cells (DG engram neurons) or ChrimsonR-negative cells (DG non-engram neurons) (fig. S6). Second, for each identified unit, the probability of spike generation evoked by stimulation of inputs from ChR2-expressing EC engram neurons served as a proxy for its functional connectivity with EC engram ensembles. This approach enabled long-term tracking of functional connectivity between EC engram inputs and spatially intermingled DG engram and non-engram neurons at single-unit resolution before and after plasticity induction.

A four-shank Neuropixels 2.0 probe was acutely inserted into the DG in combination with the optic cannula in anesthetized mice to record light-evoked spikes in DG cells (Fig. 3, D to F)(33). DG engram cells were identified by time-locked responses to 633 nm light pulses, whereas units without significant light-evoked responses were classified as non-engram cells. 473 nm light pulses were used to evoke EC engram input to DG neurons. After oSTDP stimulation, DG engram cells showed significantly decreased levels of spike probability in response to evoked EC engram inputs, whereas non-engram cells did not show such changes indicating that oSTDP specifically weakened the functional connectivity of EC-DG engram ensembles (Fig. 3, G to I). This reduction in spike generation was not mediated by changes in intrinsic excitability (Fig. 3I). Together, these data indicate that oSTDP successfully enabled a targeted weakening of EC-DG engram synapses both *in vitro* and *in vivo*.

### Memory impairment induced by synapse-specific depotentiation of EC-DG engram synapses

Finally, we tested whether the learning-induced strengthening of functional connectivity between EC-DG engram cells serves a causal role in memory expression. Optic cannulas were bilaterally implanted to the DG in Arc-CreER^T2^ mice expressing ChrimsonR-tdTomato in DG engram cells and ChR2-EYFP in EC engram cells (Fig. 4, A and B). Freezing level was measured (retrieval 1; R1) 2 days after the CFC to establish a set point of fear memory expression before *in vivo* oSTDP stimulation. Mice were subsequently moved to a neutral context and received optogenetic stimulation for oSTDP-LTD (Fig. 4A). The next day, mice were re-exposed to the fear conditioned chamber (retrieval 2; R2) to assess changes in fear response. Mice receiving oSTDP-LTD stimulation (oSTDP-LTD group) exhibited a significantly decreased freezing level (Fig. 4, G and H), whereas the memory was intact in the group of mice that received pseudo oSTDP stimulation (Fig. 4, C to E) and in Post- and Presyn-only groups (fig. S7A and fig. S7, C to E). These results indicate that the learning-induced strengthening of EC-DG engram synapses causally supports memory.

**Fig. 4.**
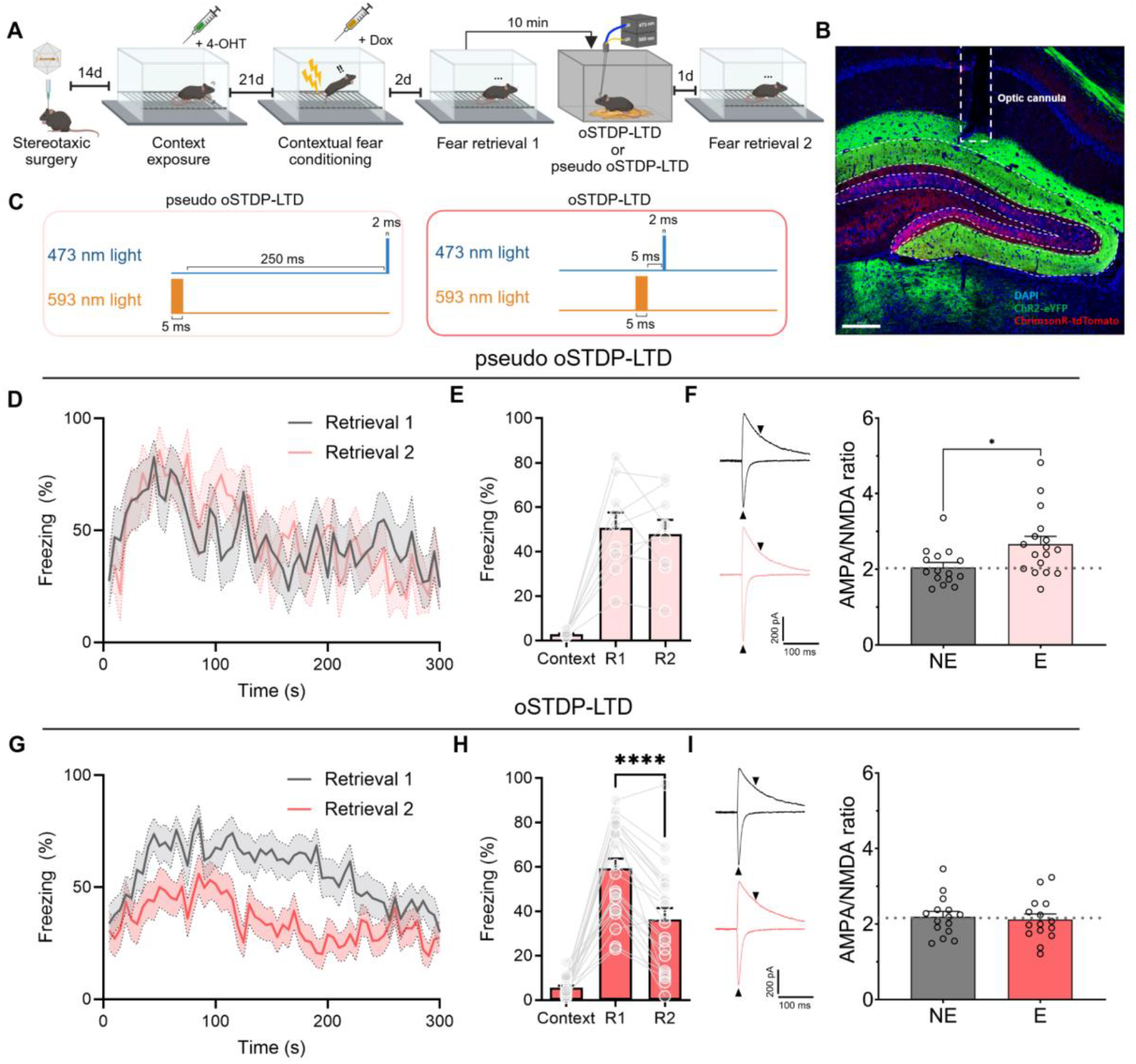
Memory impairment induced by specific manipulation of engram synapses. **(A)** Experimental schematics for memory test after *in vivo* oSTDP application to engram synapses in EC-DG circuit. **(B)** Representative image for opsin expressions in the hippocampus and optic cannula implantation. Scale bar: 200 μm. **(C)** Schematics for the description of oSTDP and pseudo oSTDP stimulation protocol used for *in vivo* experiments. **(D)** Time courses of freezing level for pseudo oSTDP-LTD group (grey, retrieval 1; light red, retrieval 2; *n* = 9 mice, *P =* 0.8641, Two-way RM ANOVA). **(E)** Mean values of freezing levels during context exposure (Context), retrieval 1 (R1) and retrieval 2 (R2) for pseudo oSTDP-LTD group (*n* = 9 mice; R1 vs R2; *P* = 0.6174, paired *t*-test). **(F)** Left, representative traces of AMPA/NMDA ratio of non-engram cell (black) and engram cell (light red) from pseudo oSTDP-LTD group. Right, AMPA/NMDA ratio of non-engram cells and engram cells from pseudo oSTDP-LTD group (grey, NE, *n* = 15 cells; light red, E, *n* = 17 cells; \**P <* 0.05; unpaired *t*-test). **(G)** Time courses of freezing level for oSTDP-LTD group (grey, retrieval 1; red, retrieval 2; *n* = 22 mice, ** *P <* 0.01, Two-way RM ANOVA). **(H)** Mean values of freezing levels during context exposure (Context), retrieval 1 (R1) and retrieval 2 (R2) for oSTDP-LTD group (*n* = 22 mice; R1 vs R2; *P* < 0.0001, paired *t*-test). **(I)** Left, representative traces of AMPA/NMDA ratio of non-engram cell (black) and engram cell (red) from oSTDP-LTD group. AMPA/NMDA ratio (grey, NE, *n* = 15 cells; red, E, *n* = 15 cells; *P =* 0.7026; unpaired t-test) for oSTDP-LTD group.

To determine whether *in vivo* oSTDP specifically alters the strength of engram synapses, we performed post-hoc *ex vivo* whole-cell recordings after oSTDP-LTD stimulation (Fig. 4I). As a control, mice received optical stimulations at intervals outside the plasticity-inducing temporal window (pseudo oSTDP-LTD; Fig. 4C). In the pseudo oSTDP-LTD group, the AMPA/NMDA ratio of EC engram synaptic inputs to DG engram cells (E-E) was significantly higher compared to non-engram cells (E-NE). However, this synaptic enhancement was absent in the oSTDP-LTD group (Fig. 4, F and I). Importantly, these oSTDP-LTD effects were memory-specific, since manipulation of synaptic ensemble encoding neutral context did not compromise fear memory expression (fig. S7B and fig. S7F).

Taken together, these results indicate that *in vivo* application of oSTDP-LTD specifically weakens EC-DG engram synapses, leading to impaired memory expression.

## Discussion

Previous attempts to reverse the learning-induced synaptic potentiation in relevant neural projection disrupted memory, demonstrating a causal link between synaptic plasticity and memory (10). Moreover, specifically disrupting the synaptic output of engram cells was sufficient to impair memory, further narrowing the locus of memory coding (12–14). Our finding that weakening of engram synapses was sufficient to disrupt memory suggests that plasticity at interregional synaptic connections between engram cells plays a crucial role in sustaining memory representation in the brain. Assigning larger synaptic weight to inputs from presynaptic engram cells to the postsynaptic engram cells may provide a mechanism by which the brain achieves memory specificity while maximizing storage capacity (14, 15).

What information is stored at the MEC/LEC-DG engram synapses remains speculative. Considering the spatial and non-spatial coding provided either individually or synergistically by MEC/LEC(34), MEC/LEC engram cells may conjunctively encode the contextual components of associative fear memory (35, 36). These streams of information may converge into DG engram cells in parallel, thereby initiating the hippocampal representation of context. This representation could be subsequently associated with valence by synaptic plasticity occurring within the downstream regions like the basolateral amygdala (37, 38). Further studies are required to define the dynamics and functions of MEC/LEC engram cells in episodic memory and to identify which components of associative fear memory are conveyed to DG engram cells.

Our data do not indicate that spike-timing dependent plasticity-like form of plasticity endogenously underlies learning (39). Rather, they show that spike-timing-dependent plasticity can be induced *in vivo* and leveraged as a tool to manipulate synaptic plasticity at functionally-defined synapses. Further investigation is required to test if this approach generalizes to other functionally-defined neuronal ensembles previously shown to display strengthened connectivity, such as like-to-like motifs in the cortex (40–44). This would help prove the causal link between neural computation and synaptic plasticity in a synapse-specific manner.

In summary, we demonstrated that enhanced functional connectivity between EC and DG engram cells causally underlies contextual fear memory. These results exemplify the causal role of the enhanced functional connectivity of neuronal ensembles in maintaining neuronal representations.

## Acknowledgments

We thank Jin-Hee Han for discussions on the manuscript. The schematic images in this study were created with BioRender.com. We used AI solely to improve the readability and language of the manuscript.

## Funding

Institute for Basic Science grant IBS-R001-D3 (B-KK)

National Research Foundation of Korea National Honor Scientist Program grant NRF-2012R1A3A1050385 (B-KK)

## Author contributions

Conceptualization: DHH, HL, JD, PP and B-KK

Methodology: YK, MC, C-HK, HL, JK, HP, CL, JYB, J-iK, D-IC, DL

Investigation: DHH, HL, JD, PP

Visualization: DHH, HL, JD

Funding acquisition: B-KK

Project administration: B-KK

Supervision: B-KK

Writing – original draft: DHH, HL, JD and B-KK

Writing – review & editing: DHH, HL, JD, PP, YK, MC, C-HK, HL, JK, HP, CL, J-YB, J-iK, D-IC, DL and B-KK

## Competing interests

Authors declare that they have no competing interests.

## Data, code, and materials availability

All data and requests for materials should be directed to the lead contact, Bong-Kiun Kaang upon reasonable request.

## Supplementary Materials

Materials and Methods Figs. S1 to S7

## Notes

### Competing Interest Statement

The authors have declared no competing interest.

