## supplementary materials for "Synaptic engram underlies memory"

**The PDF file includes:**

Materials and Methods
Figs. S1 to S7

### Materials and Methods

Mice

All experiments were carried out according to methods approved by the Institutional Animal Care and Use Committees (IACUC) at Seoul National University (SNU) and Institute for Basic Science (IBS) (IACUC #: SNU-180131-2-8, IBS-2023-038). Either heterozygous or homozygous Arc-CreERT2 (Jackson Labs; stock #021881) male mice were crossed to C57BL/6J female mice. Heterozygous Arc-CreERT2 mice and wildtype littermates of both sexes were used for experiments, aged 8–16 weeks at the time of experiment. C57BL/6J mice purchased from DBL (South Korea) were used for experiments that do not require Arc-CreERT2 based engram tagging. Mice were group-housed (maximum five mice per cage) unless otherwise mentioned and maintained on a 12 h light–dark cycle with ad libitum access to food and water.

Engram labeling systems

Stocks of 4-Hydroxytamoxifen (4-OHT; Sigma, Cat #H6278 / HelloBio, Cat #HB6040) were prepared by dissolving it in ethanol at 20 mg $ml^{-1}$ and stored at -20℃ for up to 2 months. On the day of usage, stock solution was thoroughly vortexed to redissolve 4-OHT in the ethanol to prevent dose changes due to occasional crystallization during storage. If there is no visible crystallization in the solution, 1:4 mixture of castor oil:sunflower seed oil (Sigma, Cat #S 259853 and S5007) was added to make the final concentration of 10 mg $ml^{-1}$. After gently inverting the solution to mix well with ethanol, the e-tube was placed in a vacuum centrifuge with its cap opened and allowed to ventilate for at least 30 min. The final working solution was used on the day of preparation. Doxycycline (Dox) was dissolved in saline at 5 mg $ml^{-1}$ and stored at -20℃ for up to 3 months.

4-OHT (50 mg $kg^{-1}$) or Dox (50 mg $kg^{-1}$) was administered to mice 2 hours prior to the exposure to the event (i.e. context exposure, contextual fear conditioning) by intraperitoneal (i.p.) injection under brief anesthesia by isoflurane.

Stereotaxic surgery and optic cannula implantation

Mice (3~11 weeks old at the time of surgery) were deeply anesthetized with ketamine/xylazine solution and positioned on a stereotaxic apparatus (Stoelting Co.). A long-tapered glass capillary (WPI, #504949) was prepared using the SU-P97 Puller (WPI) and the tip of the glass micropipette was cut immediately before loading of the mineral oil and virus. All virus injections were made using Nanoliter 2010 (WPI) at a 0.1 µl $min^{-1}$ rate into target regions. The tip of the glass was positioned 0.1mm below the target coordinate for 2 minutes before the injection. After injection, the glass stayed in place for an additional 7 min and was withdrawn slowly.

High titer (>$5x10^{12}$ genome copies (GC) $ml^{-1}$) of serotype 9 adeno-associated viruses ($AAV_9s$) containing following plasmids were produced by Institute for Basic Science Virus Facility: EF1α-DIO-hChR2(H134R)-eYFP, CaMKIIα -DIO-ChrimsonR-tdTomato (45), TRE3G-ChrimsonR-tdTomato and Fos-rtTA (15). Each virus was diluted in DPBS to give a final concentration stated below. 0.5 µl of AAV9-EF1α-DIO-hChR2(H134R)-eYFP ($1x10^9$ GC $\mu l^{-1}$) was bilaterally injected to both lateral entorhinal cortex and medial entorhinal cortex, unless described otherwise. A mixture of AAVs ($8x10^8$ GC $\mu l^{-1}$ of $AAV_9$-Fos-rtTA, $3x10^9$ GC $\mu l^{-1}$ of $AAV_9$-TRE3G-ChrimsonR-tdTomato) was bilaterally injected to the DG. For electrophysiology experiments in (Fig. 3C), 0.5 μl of $AAV_9$-CaMKIIα-DIO-ChrimsonR-tdTomato ($1x10^9$ GC $\mu l^{-1}$) was bilaterally injected to the DG. For some electrophysiology experiments (Fig 1G - oEPSC measurements; fig S4F), only MEC was infused with $AAV_9$-EF1α-DIO-hChR2(H134R)-eYFP ($1x10^9$ GC $\mu l^{-1}$). For experiments in (Fig. 2, D to G), $AAV_9$-CaMKIIα-iCre ($1x10^9$ GC $\mu l^{-1}$) was further mixed with $AAV_9$-EF1α-DIO-hChR2(H134R)-eYFP to be infused in MEC.

For optogenetics behavior, optic cannulas (300 μm core diameter, 0.37 NA; Newdoon) were bilaterally implanted DV 0.4 mm above the injection site of the DG after viral injection. Instant adhesive (Loctite) was applied to secure the optic implant with the skull, followed with dental cement to prevent the exposure of the skull.

For dual-eGRASP analysis, the EC of Arc-CreERT2 was infused with a total of 0.5 μl of AAV combination including: $AAV_{1/2}$-CaMKIIα-FlpO ($1.67x10^9$ GC $\mu l^{-1}$), $AAV_{1/2}$-hSyn-fDIO-preGRASP (cyan) ($1.0x10^9$ GC $\mu l^{-1}$) and $AAV_{1/2}$-EF1α-DIO-preGRASP (yellow) ($1.6x10^9$ GC $\mu l^{-1}$). Ipsilateral DG was infused with a total 0.5 μl of a combination of AAVs including: $AAV_{1/2}$-CaMKIIα-FlpO ($3.2x10^5$ GC $\mu l^{-1}$), $AAV_{1/2}$-hSyn-fDIO-myriRFP670-p2a-postGRASP ($1.6x10^9$ GC $\mu l^{-1}$), $AAV_{1/2}$-Fos-rtTA ($1.0x10^9$ GC $\mu l^{-1}$) and $AAV_{1/2}$-TRE3G-myrmScarlet-I ($1.6x10^9$ GC $\mu l^{-1}$).

Stereotaxic coordinates for each target sites were: lateral entorhinal cortex (-3.4 mm anteroposterior (AP); ±4.4 mm mediolateral (ML); -4.2 mm dorsoventral (DV)), medial entorhinal cortex (-4.6 mm AP; ±3.5 mm ML; -3.5 mm DV), and dentate gyrus (-2.0 mm AP; ±1.35 mm ML; -2.15 mm DV (from skull surface)).

*In vitro* electrophysiology

Mice were deeply anesthetized by the i.p. injection of ketamine/xylazine solution and transcardial perfusion was performed with ice-chilled sucrose solution which contained (in mM): 210 sucrose, 3 KCl, 26 $NaHCO_3$, 1.25 $NaH_2PO_4$, 5 $MgSO_4$, 10 D-glucose, 3 sodium ascorbate and 0.5 $CaCl_2$, saturated with 95% O2 and 5% CO2. Subsequently, the brain was isolated and immediately placed in the ice-chilled sucrose solution for ~30 s. The hippocampi were rapidly isolated from the brain, transverse hippocampal slices of 350 μm were prepared using a vibratome (Leica, VT1200S). Slices were transferred to the recovery chamber filled with the artificial cerebrospinal fluid (ACSF) which contained (in mM): 124 NaCl, 3 KCl, 26 NaHCO3, 1.25 NaH2PO4, 2 MgSO4, 10 D-glucose and 2 CaCl2 (carbonated with 95% O2 and 5% CO2) for 30 min at 32-34°C. The temperature of the recovery chamber was then adjusted to 28-30°C and maintained until the recording was made.

Whole-cell recordings were conducted in the recording chamber that was constantly perfused with ACSF, to which picrotoxin (Hellobio, HB0506) 100 μM added, at 3-4 ml/min at 32°C. Borosilicate glass pipettes were used with a resistance of 3-6 MΩ. K-gluconate internal solution consisting of (in mM): 8 NaCl, 130 K-gluconate, 10 HEPES, 0.5 EGTA, 4 Mg-ATP, 0.3 $Na_3$-GTP, 5 KCl and 0.1 spermine was used for optical STDP-LTD experiments (Fig. 2D to G; Fig. 3C). K-$MeSO_3$ internal solution consisting of (in mM): 8 NaCl, 130 K-$MeSO_3$, 10 HEPES, 0.5 EGTA, 4 Mg-ATP, 0.3 $Na_3$-GTP, 5 KCl and 0.1 spermine was used in some experiments (fig. S3). $CsMeSO_3$ internal solution consisting of (in mM): 8 NaCl, 130 $CsMeSO_3$, 10 HEPES, 0.5 EGTA, 4 Mg-ATP, 0.3 $Na_3$-GTP, 5 QX-314 and 0.1 spermine was used for the measurement of AMPA/NMDA ratio and paired-pulse ratio (PPR). The pH was adjusted to 7.2-7.3 either with KOH or CsOH and osmolarity was set to 285-290 mOsm/l. Signals were obtained with Multiclamp 700B amplifier (Molecular Devices) and digitized with Digidata 1550 (Molecular Devices, US) at a sampling rate of 20 kHz (filtered at 2 kHz).

TdTomato positive neurons in DG were visualized by Axio Examiner.D1 microscopy (40x, ZEISS, US). Excitation light was illuminated by CoolLED pE340 fura (CoolLED, UK) and a real-time image was delivered to the computer through a SCMOS digital camera. Neurons were voltage clamped at -70 mV and spontaneous responses were monitored for at least 5 min after the whole cell configuration was established to stabilize the effect of internal solution influx.

In experiments using optogenetics for in vitro electrophysiology, we only used slices with successful virus expression, which was selected by group-blinded experimenters based on opsin expression intensities of the DG granule cells and axons in the perforant pathway. ChR2 expressed in axons of EC cells was activated by 473 nm light (0.5~2mW, 1 ms) shining, which did not evoke a substantial ChrimsonR opening under our experimental conditions. Engram cells were identified with a substantial amount of ChrimsonR-mediated inward currents and tdTomato+. Synaptic inputs were activated with an inter-stimulation interval of 6s.

For oSTDP experiments in (Fig. 3C), hippocampal slices were prepared within 30 min after memory retrieval for oSTDP in vitro experiments. After obtaining at least 5 min of stable baseline amplitude of monosynaptic excitatory postsynaptic currents (EPSCs), 5 min pairing of EPSCs and postsynaptic depolarization was given to the cell. Optical STDP protocol consisted of alternating light stimulation (2 Hz, 600 pairing, 5 ms interval between 473nm light (0.5~2 mW, 1 ms duration) and 593 nm light (4~5 mW, 5ms). Stimulation with 593-nm light consistently evoked substantial depolarization but action potentials were not consistently evoked. The intensity of 473-nm light was adjusted to remain subthreshold during oSTDP induction. For the analysis shown in (Fig. 2G), cells were pooled regardless of whether 593-nm light stimulation evoked action potentials during oSTDP induction as long as substantial depolarization by 593-nm light stimulation was observed. For non-specific synaptic input activation in (fig. S3), the activation of afferent MEC synaptic inputs was achieved by stimulating electrodes placed in the two-third part of the molecular layer and APs were evoked by current injection (300~400 pA, 5 ms).

For AMPA/NMDA ratio, cells were clamped at a holding potential of -70 mV to measure the peak of AMPAR-mediated synaptic transmission. NMDAR-currents were estimated at 50 ms after the stimulation onset at +40 mV of holding potential. Averages of at least 5 consecutive responses obtained at these holding potentials were used for the ratio. Holding voltage of 0 mV was used for the quality control of AMPA/NMDA ratio measurement for each cell. For PPR, cells were voltage clamped at -70 mV and 5 consecutive paired stimulation with interval varying from 50 ~ 200 ms was given to estimate presynaptic release probability. For *ex vivo* experiments in (Fig. 4, F and I), group-blinded experimenters determined whether the virus targeting in subject was successful by checking the ChR2-EYFP expression pattern of MEC and LEC projections and used for the experiment only if at least three projections out of total four projections were judged to be successful. Further, only hippocampal slices with visible track of the optic cannula were used.

*In vivo* electrophysiology

*Surgery and behavior*

Mice used for in vivo electrophysiological recordings were infused with AAV9-CaMKIIα-DIO-ChrimsonR and AAV9-EF1α-DIO-ChR2-EYFP into the right hemisphere DG and EC, respectively. After 2 weeks of recovery, mice were injected with 4-OHT 2 hours before context exposure, as described in the "Engram labeling systems" section. Two weeks later, mice were injected with 4-OHT (50 mg $kg^{-1}$) and re-exposed to the same context, where they received three foot shocks (2 s duration, 0.75 mA). One week later, surgeries for optic cannula implantation and Neuropixels recordings were conducted.

On the first day, mice were deeply anesthetized by intraperitoneal injection of a ketamine/xylazine solution and placed in a stereotaxic apparatus (Stoelting Co.). An optic cannula (200 μm core diameter, 0.37 NA; Newdoon) was implanted in the right hemisphere, with the cannula tip positioned 0.35 mm dorsal to the DG injection site. A stainless-steel head bar was affixed to the skull using light-cured dental composite and dental acrylic to minimize head movement during Neuropixels recordings. The skull was then covered with Kwik-Cast to protect the exposed surface until the second surgery.

On the second day, mice were anesthetized with isoflurane (1–2%) and head-fixed in a stereotaxic frame (Narishige, Japan) using the previously implanted head bar. Body temperature was maintained at 37°C using a heating pad with a DC temperature control system (FHC Inc., Bowdoin, ME, USA). A stainless-steel screw for external reference was implanted in the left hemisphere at a position that would not interfere with the trajectory of the Neuropixels probe. A circular craniotomy with a diameter of 1.5 mm was made over the target site using a microdrill at AP −1.9 mm and ML -3.15 mm. The cortical surface was continuously covered with PBS to prevent drying until probe insertion.

Recordings were performed using four-shank Neuropixels 2.0 probes. Before insertion, the tips of all four shanks were coated with DiD (Vybrant™ DiD Cell-Labeling Solution, V22887) to allow subsequent histological verification of the probe tracks. The probe was mounted on a four-axis micromanipulator (UNIT Junior 4 axes with SM10 Compact control box, Luigs & Neumann, 210-100 000 0080). Because the optic cannula was implanted vertically, the Neuropixels probe was inserted at a 40° angle from the vertical axis to avoid collision with the cannula. After the DiD coating, the probe was slowly inserted through the craniotomy at a speed of 2–8 μm/s and advanced until the tips of the shanks were positioned at AP -2.0 mm, ML -1.35 mm, and DV -2.15 mm. After reaching the target depth, the probe was advanced an additional 100 μm and then retracted by 100 μm to minimize tissue compression. To reduce recording drift, the probe was allowed to stabilize for 1 hour after insertion before data acquisition.

*Experimental procedure*

Each recording session consisted of three phases: pre-oSTDP optogenetic stimulation, optical STDP (oSTDP), and post-oSTDP optogenetic stimulation. The oSTDP phase was performed using the same stimulation protocol described below (see ‘optogenetics behavior’). Laser stimulation was delivered using Omicron LuxX diode lasers. Wavelengths of 473 nm and 633 nm were used for ChR2 and ChrimsonR activation, respectively.

Each optogenetic stimulation phase consisted of separate 473 nm and 633 nm light stimulation blocks. 633 nm light stimulation was performed to determine whether recorded units were ChrimsonR-positive, and therefore only two light intensities, 20 and 40 mW, were used. In contrast, ChR2 stimulation was performed using a broad range of light intensities from 0.1 to 20 mW to sensitively assess changes in light-evoked responses induced by oSTDP. This range was chosen to avoid floor and ceiling effects in detecting oSTDP-induced changes: low-intensity stimulation may fail to evoke spikes even in the presence of oSTDP-induced changes, whereas high-intensity stimulation may reliably evoke spikes regardless of oSTDP-induced changes, potentially masking subtle changes in response probability.

The duration of light stimulation was matched to the oSTDP protocol: 473 nm light was delivered for 2 ms, and 633 nm light was delivered for 5 ms. For mice showing particularly strong viral expression, shorter stimulation durations, ranging from 0.2 to 1 ms, were used to avoid saturated responses. Each light intensity was repeated 10 times. Trials were presented in an order randomized across light intensities, with an inter-stimulus interval of 12 s. The pre-oSTDP and post-oSTDP optogenetic stimulation phases used the same light intensities and stimulation durations. The entire recording session, including all three phases, lasted approximately 2 h.

*Data acquisition*

Extracellular recordings were performed using a four-shank Neuropixels 2.0 probe, National Instruments PXI hardware, and SpikeGLX software. Recordings were made from all four shanks, with 96 recording channels per shank (384 channels in total) positioned near the shank tips and spanning 720 µm along each shank. Neuropixels data were acquired at 30 kHz. Optogenetic stimulation was controlled by Arduino-generated TTL pulses sent to the laser controller, while duplicate TTL pulses were recorded by the NI-DAQ at 10,593.2 Hz to mark stimulation timing. A 1-Hz synchronization signal generated by the Neuropixels headstage was recorded by the NI-DAQ throughout each session and used for offline alignment of the electrophysiology data with optogenetic stimulation timing.

*Data analysis*

Neuropixels recordings were preprocessed using CatGT. Individual trials were concatenated and filtered with a 12th-order Butterworth band-pass filter from 1,000 to 10,000 Hz to reduce low-frequency light-induced artifacts. To remove transient light artifacts, 1-ms segments (30 samples) were masked around each stimulation onset and offset. Global demultiplexed common average referencing was then applied (option -glbdmx).

Spike sorting was performed using the Python implementation of Kilosort 4.0 (46). By default, Kilosort classifies clusters with a refractory-period contamination percentage (the relative increase in spikes occurring within the refractory period compared with baseline firing) greater than 20% as multi-unit activity (MUA). This default GOOD/MUA classification was not used as a unit-selection criterion because some dentate gyrus units were nearly silent during baseline periods but exhibited reliable short-latency firing after light stimulation, which could result in inflated contamination estimates despite clear light-evoked responses.

Instead, units were selected using stringent waveform- and amplitude-based criteria. Only units with a waveform amplitude signal-to-noise ratio greater than 3 were considered. For each candidate unit, a 1-ms template waveform was extracted from the peak channel and the four nearest channels. This five-channel, 1-ms template was then shifted sample by sample across the entire recording, and its correlation with the corresponding signal segment was calculated at each time point. Events were retained only if the waveform–template correlation was $\geq 0.9$ and the spike amplitude was within $\pm 20\%$ of the unit′s mean spike amplitude. Sessions with an average drift greater than one Neuropixels channel spacing (20 µm) were excluded from analysis. During the optogenetic stimulation phase, units were classified as DG engram units if they generated a spike in at least 10% of all 633-nm stimulation trials, corresponding to a response fidelity of $\geq 0.1$. Approximately 1% of sorted units met these criteria and were included in the analysis.

Optogenetics behavior

All mice were introduced for behavioral experiments after a minimum two weeks of recovery after surgery. For context labeling, 4-OHT (50 mg $kg^{-1}$) was injected by intraperitoneal injection during brief anesthesia by isoflurane 2 hours prior to the context exposure. Mice were exposed to a square chamber with a metal grid (MED associates) for 300 s without foot shock. Mice did not show freezing behaviors during the initial exposure to the context. For contextual fear conditioning, either Dox or 4-OHT (50 mg $kg^{-1}$) was injected by intraperitoneal injection 2 hours before the conditioning (see, experimental schematics in each figure and method 'Engram labeling systems' section). Mice were reintroduced into context where three 0.75 mA shocks of 2 s duration were delivered within a total 300 s of exploration. For testing fear memory retention, mice were subjected to 300 s retrieval session in context 2 days (or 1 day for fig. S7F) post training.

For optical STDP stimulation, optic cannula implanted to mice were tightly fit with patch cord (Newdoon) that was connected to 473/593 nm laser via light spectrum mixer (Doric) and rotary joint (Doric) in a new cage with bedding inside a distinct room with dim light. Mice were allowed to freely explore the context for 10 min and were delivered with optical STDP stimulation consisting of: 593 nm light (15 mW at fiber tip) of 5 ms pulse width followed by 473 nm light (0.8 ~ 10 mW at fiber tip) of 2 ms pulse width, with 5 ms interval. The stimulation was delivered for 15 min (2 Hz). Mice were immediately detached from the patch cords after the stimulation and were returned to their home cages. Subsequent testing sessions were conducted a day after the initial retrieval session. For in vivo oSTDP-LTD experiments, we only included mice in which viral expression was confirmed in at least three out of the four perforant paths (MEC and LEC in both hemispheres) and both dentate gyri were well infected. All exclusion was done in a group blinded manner.

For DG engram reactivation, trained mice were attached to the patch cord in a distinct room mentioned above. Mice remained in their homecages for at least 20 minutes and were relocated to an acryl box for optical stimulation, which consisted of four 3-minute epochs, with the first and third epochs serving as the light-off epochs, and the second and fourth epochs as the light-on epochs. During the light-on epochs, mice received light stimulation (15 ms pulse width, 20 Hz) for the entire 3-minute duration. At the end of 12 minutes, mice were immediately detached from the patch cords and returned to their home cages. Freezing level was manually counted by a group-blinded experimenter. We only included mice with verified viral expression in the targeted regions.

Sample preparation and confocal imaging

Mice were anesthetized with ketamine/xylazine solution for transcardial perfusion. The perfused brains were stored in 4% PFA in PBS overnight at 4°C and then dehydrated in 30% sucrose for 2 days. Brain slices of 40~50 µm (40 µm for IHC, 50 µm for dual-eGRASP analysis) were prepared using cryostat (Leica). Sections were mounted in a Vectashield mounting medium (Vector Laboratories). EC and DG were imaged in Z-stack with Leica SP8 confocal microscope with a 20x objective, and for dual-eGRASP analysis, DG dendrites were imaged in Z-stack with a 63x objective and distilled water immersion.

Image analysis for dual-eGRASP

Analysis of dual-eGRASP and 3D reconstruction of DG dendrites were performed using IMARIS (Bitplane, Zurich, Switzerland) software as previously described (15). Each mScarlet-I-positive or iRFP670-positive dendrite was manually marked as a filament, while hiding other fluorescent signals. The identity of postsynaptic neurons was determined by the fluorescent protein expressed through the membrane where mScarlet-I and iRFP670 co-expressing cells were counted as DG engram neurons and iRFP only cells were counted as DG non-engram neurons. Each cyan or yellow eGRASP signal was marked as cyan or yellow spheres through IMARIS automatic detection. Yellow only or yellow + cyan eGRASP signal was counted as synapse originating from EC engram cells, while cyan only eGRASP signal was counted as synapse originating from EC non-engram cells. Cyan and yellow eGRASP puncta on dendrites were manually counted. The identity of each dendrite was blinded during the analysis to exclude any bias. Spine density data were normalized within one confocal image based on the raw value of GRASP density of iRFP dendrite for further statistical analysis.

Immunohistochemistry (IHC)

Brain slices were washed with 1 ml PBS for a minute 3 times at room temperature (RT), 120rpm. 400 µl of blocking solution (3% goat serum, 0.3% triton X-100 in PBS) was added to each well for 1 hour, 80 rpm. Each well contained no more than 4 slices. Primary antibody was treated at the concentration of 1:1000 and incubated for 24 hours while being shaken. The next day, slices were washed with 0.2% PBST for 3 times 10 minutes each at 120 rpm, RT. Secondary antibody was treated at the concentration of 1:400 for 2 hours, 80 rpm. Finally, sections were washed with 1 ml PBST 3 times for 10 minutes at 120 rpm, RT. DAPI was treated for 5 minutes, 80 rpm, RT. Sections were washed with 1 ml PBS for 3 times for 5 minutes 120 rpm, RT. c-Fos was stained with rabbit anti- c-Fos (1:1000, 226 003, SySy) and anti-rabbit Alexa-488 (1:500). Cell counting was manually done with Imaris (Bitplane) by a blinded experimenter.

Statistics

Data were analyzed using Prism software. Unpaired t-test, paired t-test, Wilcoxon matched-pairs signed rank test, one-sample t-test, Mann-Whitney two-tailed test and Tukey's multiple comparison test either after RM one-way ANOVA or RM two-way ANOVA were used to test for statistical significance when applicable. The exact value of n and statistical significance are reported in each figure legend. A mouse in oSTDP-LTD behavior group was excluded from (Fig. 4, G and H) based on an outlier test and exclusion did not change the statistical significance. Data are presented as mean ± standard error of the mean (s.e.m.).

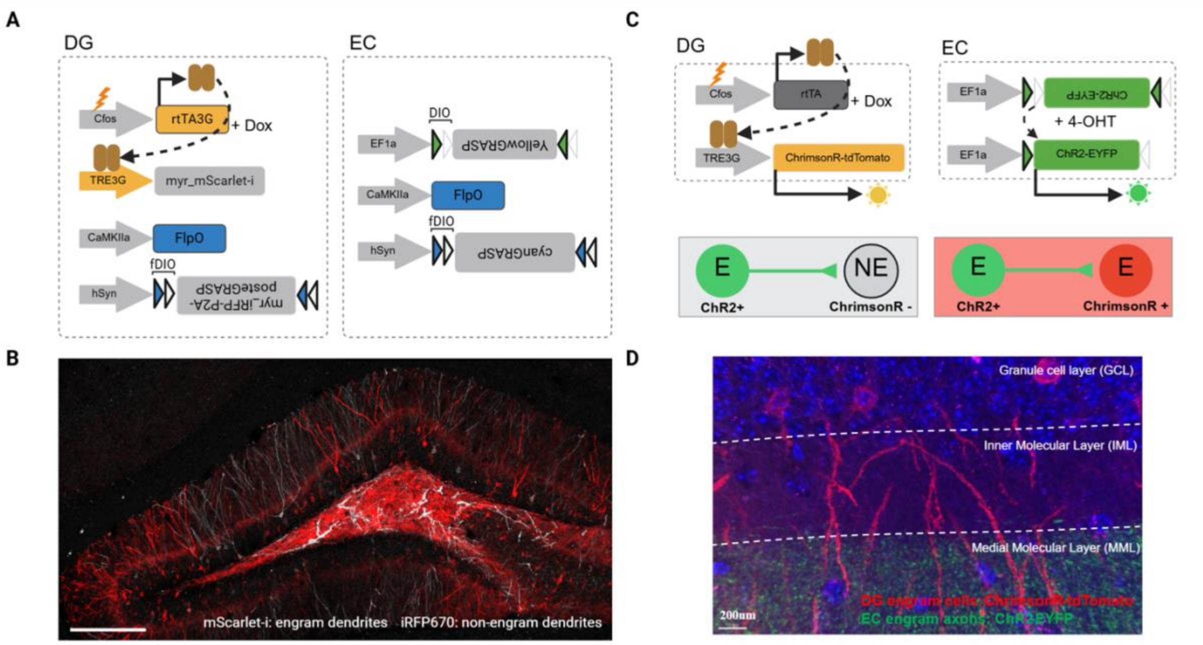


**Fig. S1. Recruitments of engram cells in DG and EC following contextual fear memory formation.**

(A) Schematics showing viral combinations used for dual-eGRASP experiments in (Fig. 1, A to D). (B) A representative image of hippocampal DG slice used for the quantification of the densities of synaptic connections between EC and DG engram cells. Scale bar: 200 µm. (C) Schematics showing viral combinations used for whole cell recordings in (Fig. 1, E to G). (D) Representative hippocampal image showing an example DG engram cell (tdTomato+) and axons of EC engram cells (EYFP+). Scale bar: 200 µm.

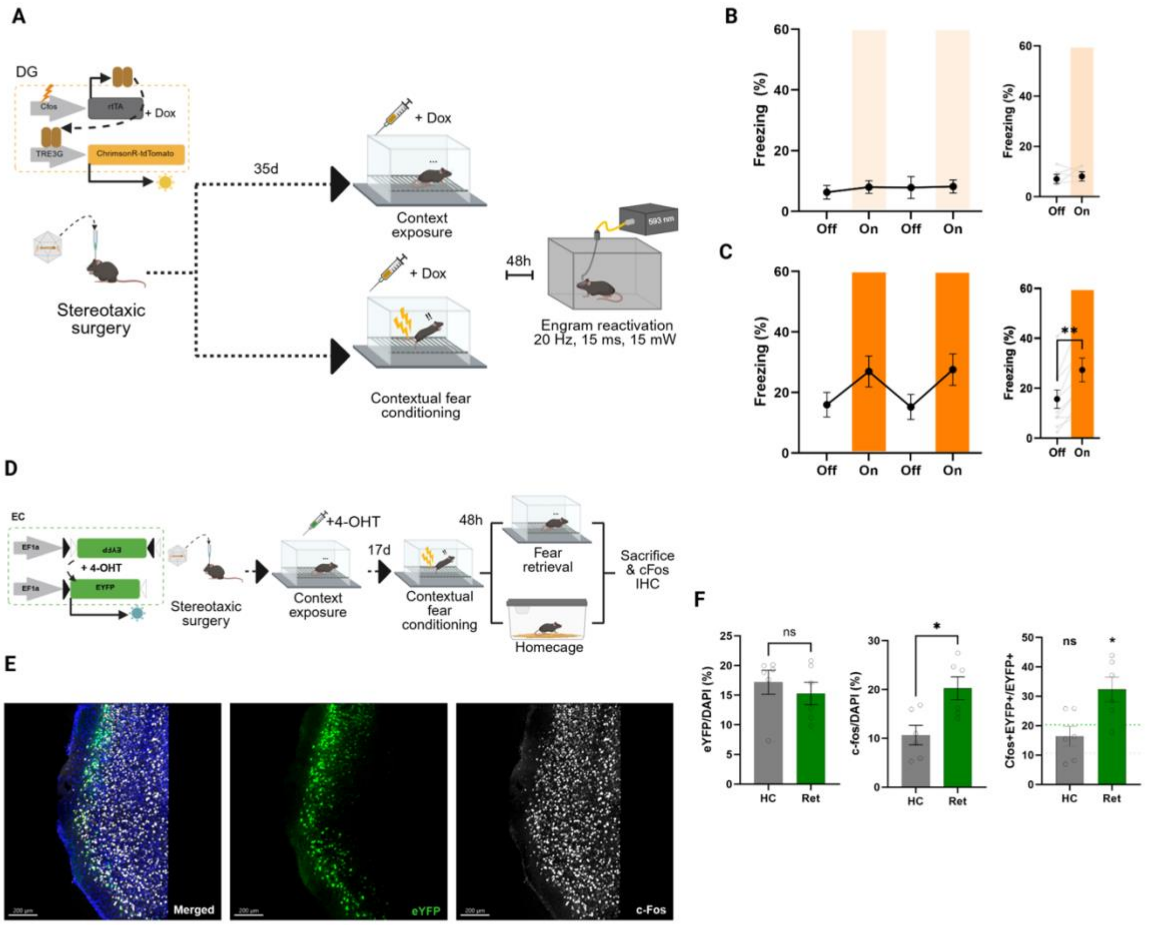


**Fig. S2. Recruitments of engram cells in DG and EC following contextual fear memory formation.**

(A) Experimental schematics for DG engram reactivation. (B) Left, freezing level of context exposure group during each light off/on epochs (n = 6 mice). Right, comparison of freezing levels of off/on epochs (n = 6 mice; P = 0.6875, Wilcoxon matched-pairs signed rank test). (C) Same as with (B) but for fear conditioned group (n = 10 mice; **P < 0.01, Wilcoxon matched-pairs signed rank test). (D) Experimental schematics for the measurement of EC engram cells. (E) Representative images showing context-exposure activated neurons (EYFP+) and fear-recall activated neurons (iRFP+). (F) Left, percentages of context-exposure activated neurons in EC (grey, homecage, HC; green, fear retrieval, Ret). Middle, percentages of cFos-positive neurons (grey, homecage, n = 6 mice; green, fear retrieval, n = 6; *P < 0.05, Welch's t test). Right, reactivation ratios of context-exposure activated neurons during either in the home cage (grey, n = 6 mice; P = 0.1787, Welch's t test compared to the chance level) or fear retrieval session (green, n = 6 mice; P = 0.0360, Welch's t test compared to the chance level). The dotted line indicates the chance level calculated from the percentages of cFos-positive neurons.

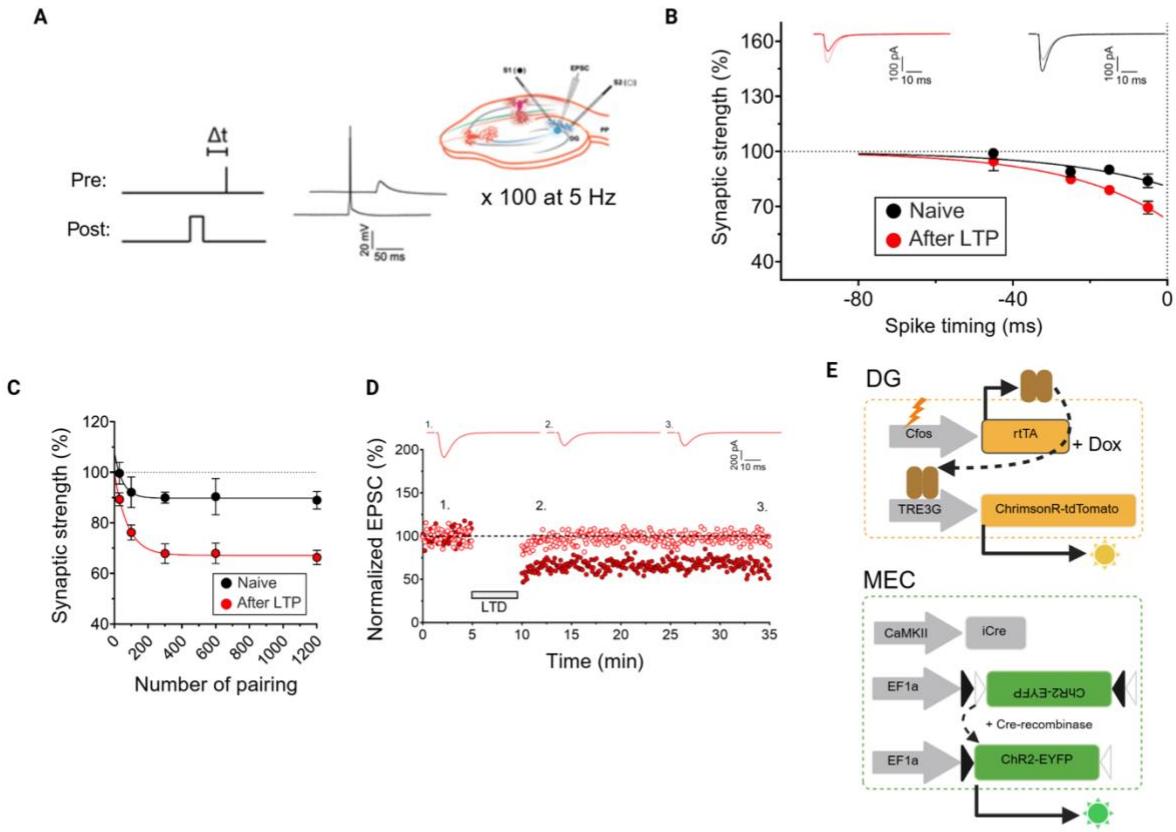


**Fig. S3. Characterization of STDP rules in MEC-DG projection.**

(A) Example trace for STDP inducing stimulation and an illustration for experimental conditions. (B) A plot showing a temporal window of spike-timing-dependent plasticity (STDP) at medial perforant path to dentate gyrus granule cells. Changes in the synaptic strength were plotted against the EPSC-action potential (AP) delays; black and red dots represent values obtained from naive slices and slices that underwent LTP-inducing stimulation, respectively (+45 ms (n = 5/5), +25 ms (n = 6/6), +15 ms (n = 6/7), +5 ms (n = 13/8), -5 ms (n = 8/12), -15 ms (n = 6/8), -25 ms (n = 6/6), -45 ms (n = 4/6), numbers in parentheses indicate number of neurons in naive slice/prior LTP slice); negative value in x-axis indicates that EPSC followed AP. Inset displays representative traces before and after STDP-inducing stimulation; light red and black traces before STDP in prior LTP/naive condition; red and grey after STDP in prior LTP/naive condition. (C) A plot showing resultant changes in synaptic strengths against the number of pairing during STDP-inducing stimulation. Synaptic strengths are obtained by averaging the amplitudes of the last 5 sweeps of recordings that followed STDP-LTD stimulation (black, naive slice; red, after LTP; 30 pairings (n = 3/4), 100 pairings (n = 5/9), 300 pairings (n = 3/5), 600 pairings (n = 3/4), 1200 pairings (n = 3/4), numbers in parentheses indicate number of neurons in naive slice/after LTP slice). (D) Representative trace for STDP-LTD inducing stimulation. Filled red dots represent EPSC values of test inputs where STDP-LTD stimulation was given and empty red dots represent control inputs. EPSC amplitudes are normalized to baseline level. (E) Schematics showing viral combinations used for whole cell recordings in (Fig. 2, C to G).

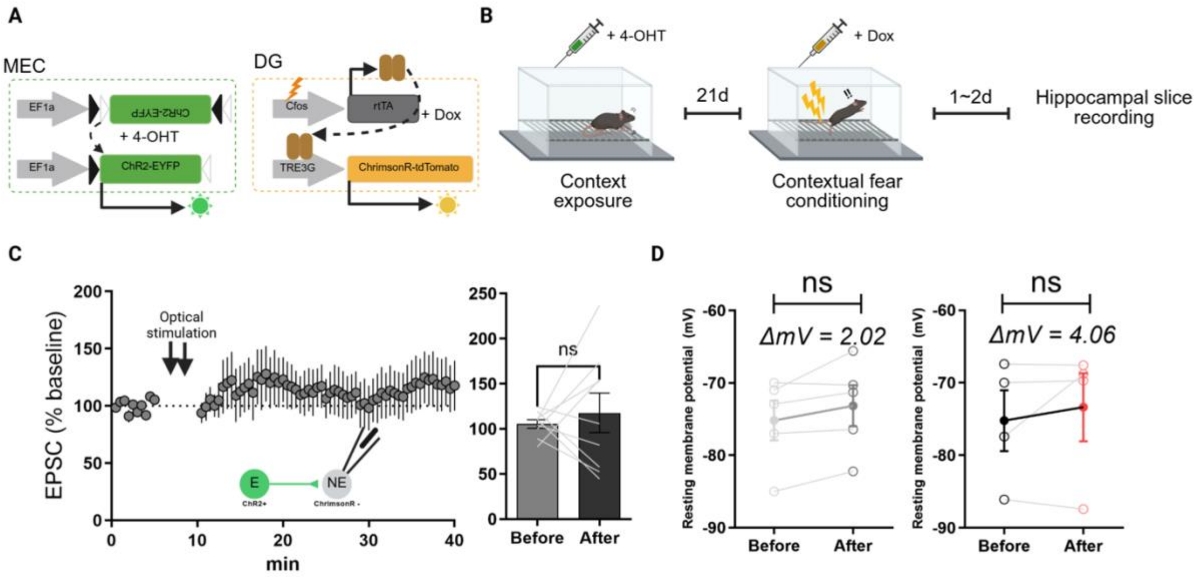


**Fig S4. Recruitments of engram cells in DG and EC following contextual fear memory formation.**

(A) Schematics showing viral combinations used for whole cell recordings in (fig. S5). (B) Experimental scheme for oSTDP-LTD validation in slice recording. (C) Left, a time-course of MEC engram input-evoked oEPSCs amplitude of DG non-engram cells after oSTDP-LTD. Right, the average amplitude of the last 5 sweeps of EPSCs during baseline period and the last 5 sweeps of EPSCs 30 min after optical stimulation (n = 9 cells, ns; paired t-test). (D) Resting membrane potential of DG engram cells and non-engram cells before/after oSTDP stimulation (left, non-engram cells, n = 5 cells, ns; engram cells, n = 4 cells, ns; paired t-test).

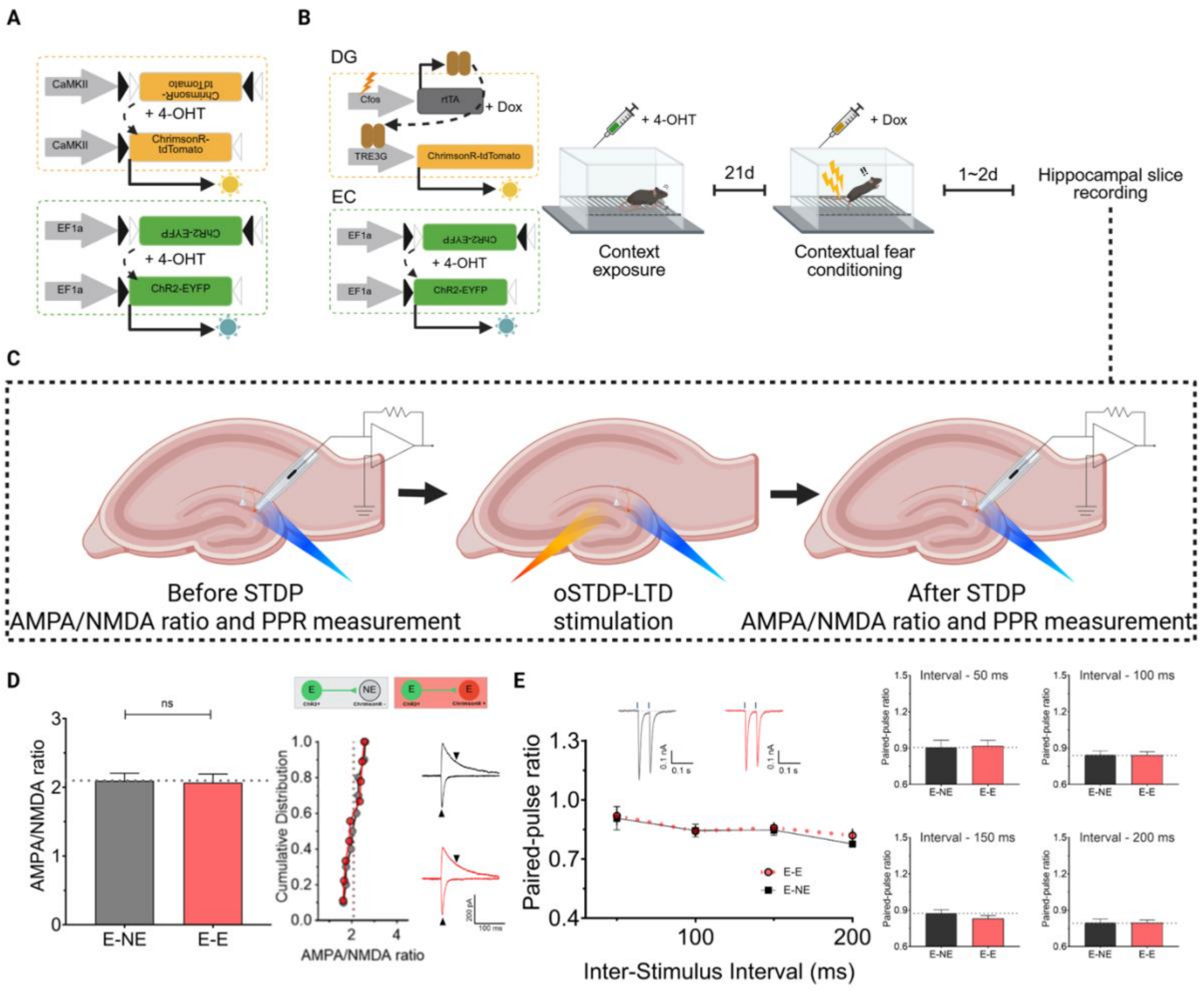


**Fig S5. AMPA/NMDA ratio and PPR of DG engram cells following *in vitro* STDP-LTD.**

(A) Schematics showing viral combinations used for whole cell recordings in (Fig. 3, A to C). (B) Experimental scheme for oSTDP-LTD validation in slice recording. (C) Synaptic inputs from EC engram cells were evoked by ChR2 activation of EC engram axon terminals while recording DG engram or non-engram cells' responses under whole-cell recording configuration. At least 10 min after oSTDP-LTD stimulation given to the hippocampal slice, DG engram or non-engram cells were recorded to examine pre-/postsynaptic expression of oSTDP-LTD mediated LTD. (D) Left, AMPA/NMDA ratio of E-NE or E-E synapses after oSTDP-LTD stimulation *in vitro* (E-NE, grey, n = 10; E-E, red, n = 9 cells; P = 0.8592, unpaired t-test). Middle: cumulative distribution of the same data set of AMPA/NMDA ratio of E-NE versus E-E. Dotted lines indicate the mean values for each group. Right, example traces of AMPA-/NMDA receptor-mediated currents of a DG engram or non-engram cell. (E) Left, paired-pulse ratios (PPRs) for E-NE (n = 10 cells) and E-E (n = 9 cells) in 50 ms, 100 ms, 150 ms and 200 ms after oSTDP-LTD stimulation *in vitro*. Insets show an example trace of E-NE and E-E to paired stimuli of inter-stimulus interval of 50 ms. Right: bar graph displaying mean values of PPRs for each interval. 50 ms, P = 0.8744; 100 ms, P = 0.9443; 150 ms, P = 0.28175; 200 ms, P = 0.9769, unpaired t-test.

**A.** DG engram cells

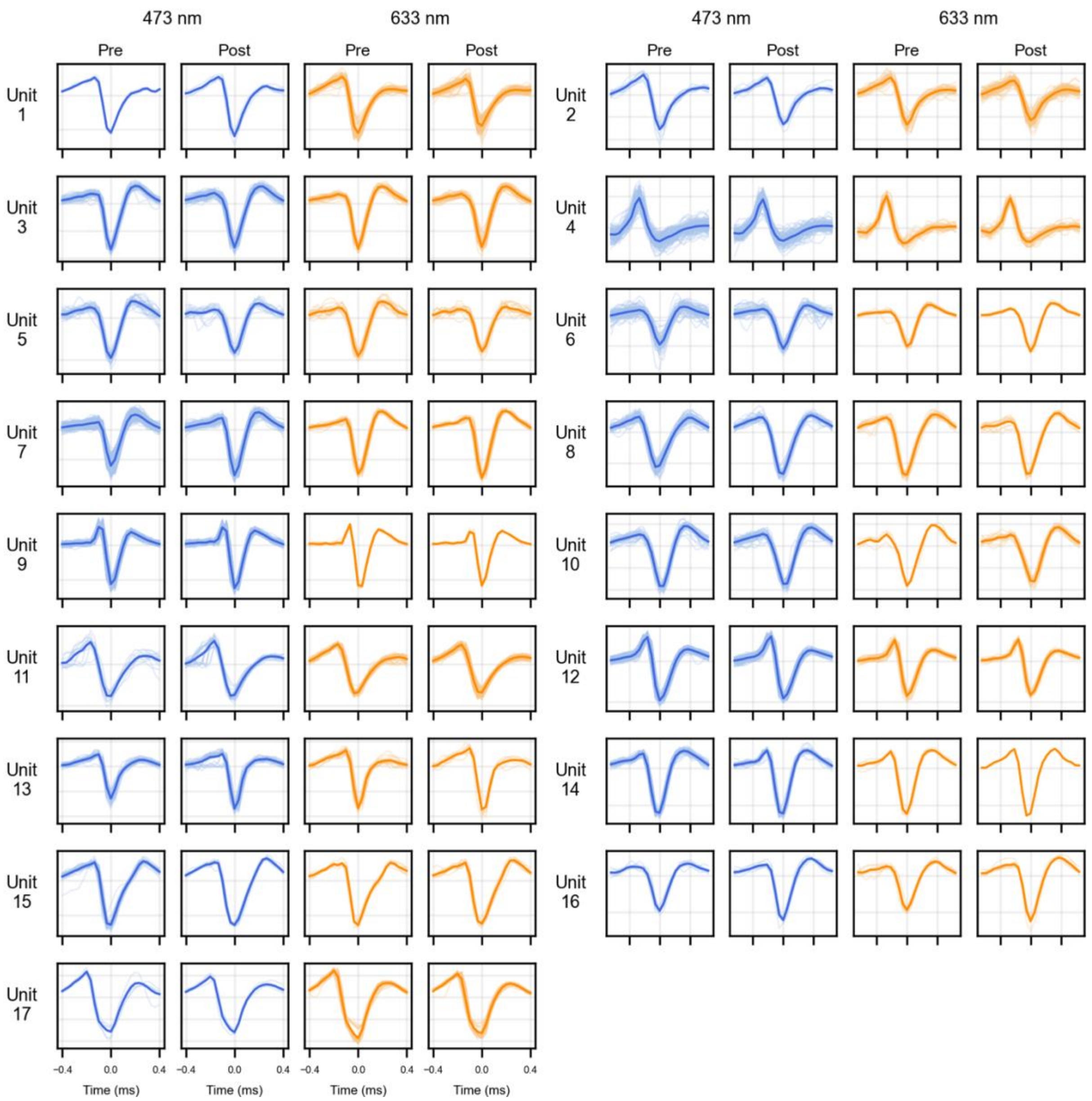

**B.** DG non-engram cells

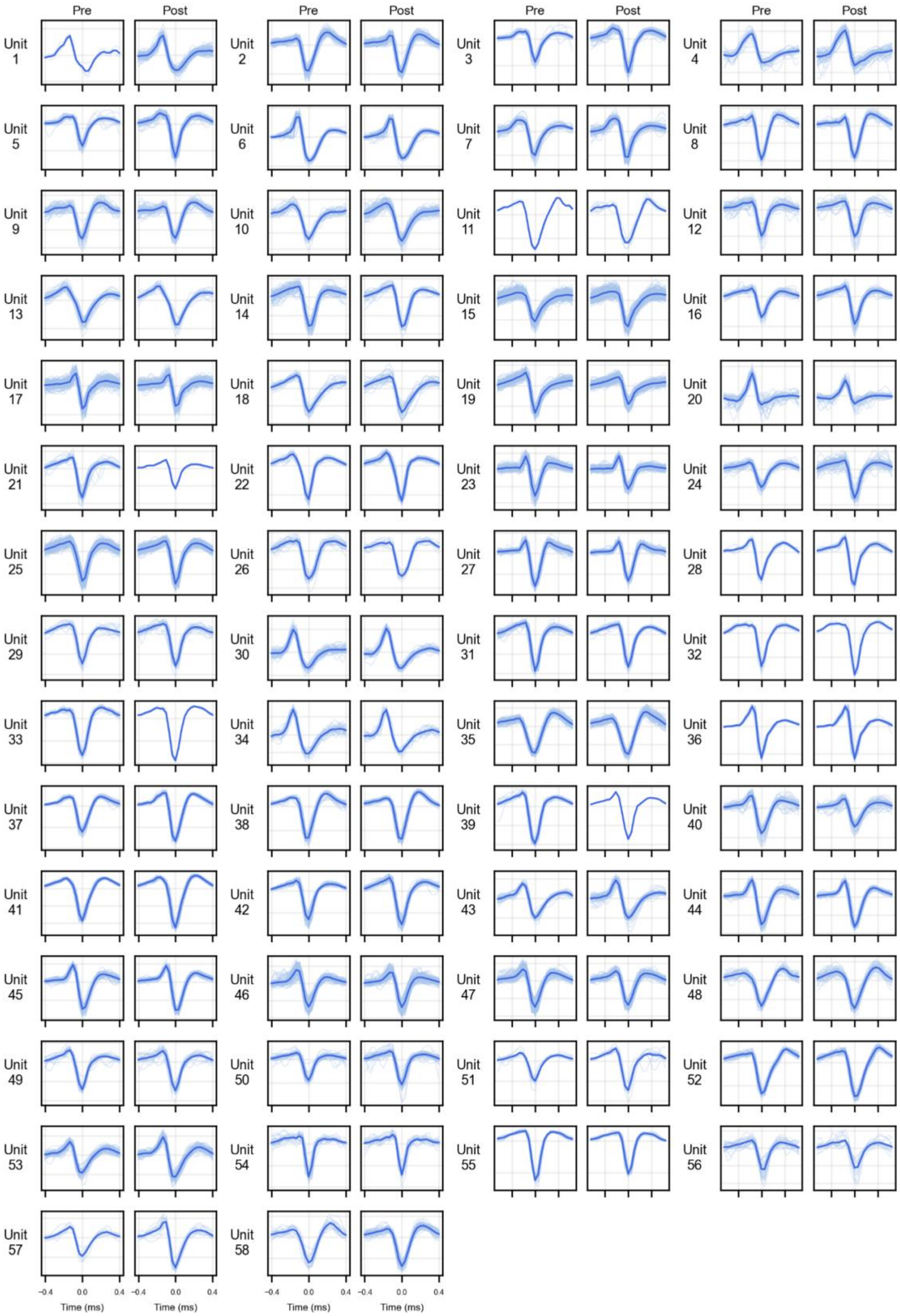

**Fig S6. Individual light-evoked waveforms per unit.**

(A) Light-evoked spike waveforms from DG engram cells during the Pre- and Post-oSTDP optogenetic stimulation phases. Thin traces represent individual spikes, and the thick trace represents their mean waveform. The four panels for each unit show responses to 473-nm and 633-nm stimulation before and after oSTDP (blue, 473 nm-evoked; orange, 633 nm-evoked). (B) Light-evoked spike waveforms from DG non-engram cells during the Pre- and Post-oSTDP optogenetic stimulation phases. Only 473-nm-evoked responses are shown because DG non-engram units were defined by the absence of a response to 633-nm stimulation.

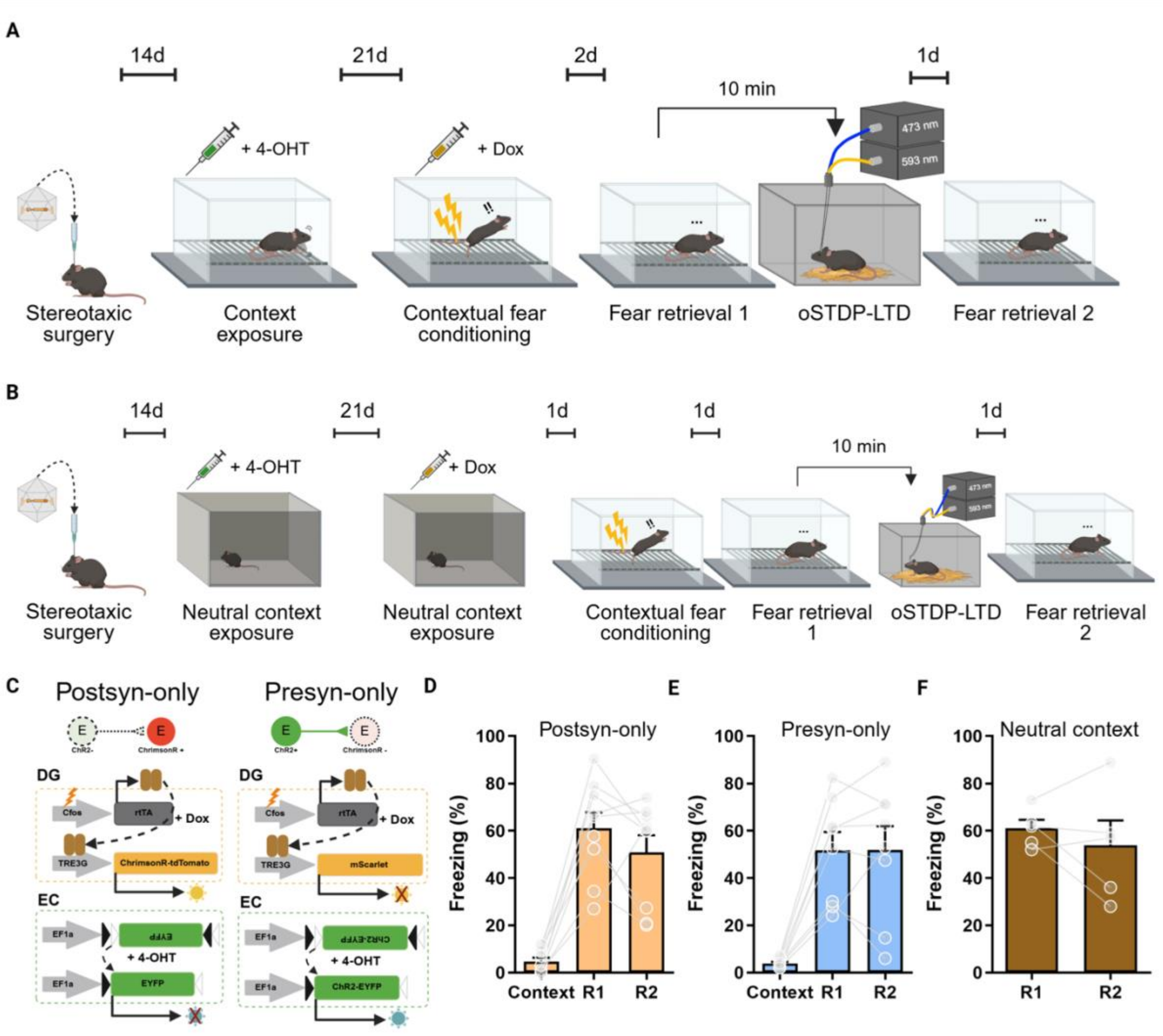


**Fig S7. No significant changes of memory expression by control stimulations or manipulation of synapses encoding other memory.**

(A) Behavioral schedule for control groups of in vivo oSTDP-LTD. (B) Behavioral schedule for labeling engram cells encoding novel context and manipulation via in vivo oSTDP-LTD. Same AAVs combinations used for the oSTDP-LTD group in (Fig. 4) were injected. (C) AAVs combination for control groups. (D) Mice that did not express ChR2 in EC engram cells (Postsyn-only, tangerine, n = 9 mice; P = 0.2071, paired t-test; R1 vs R2) did not show changes in the freezing level. (E) Mice that did not express ChrimsonR in DG engram cells (Presyn-only, blue, n = 8 mice; P = 0.9776, paired t-test; R1 vs R2) did not show changes in the freezing level. (F) A cohort of mice of which engram cells were labeled in a neutral context did not show changes in the freezing level (n = 5 mice; P = 0.4509, paired t-test; R1 vs R2).
